# Synergistic control of axon regeneration and functional recovery by *let-7* miRNA and Insulin signalling (IIs) pathways

**DOI:** 10.1101/2024.08.03.606500

**Authors:** Sruthy Ravivarma, Sibaram Behera, Dipanjan Roy, Anindya Ghosh-Roy

## Abstract

The capability of neurons to regenerate after injury becomes poor in adulthood. Previous studies indicated that loss of either *let-7* miRNA or components of Insulin signalling (IIs) can overcome the age-related decline in axon regeneration in *C. elegans*. In this study, we wanted to understand the relationship between these two pathways in axon regeneration. We found that the simultaneous removal of *let-7* and the gene for insulin receptor *daf-2* synergistically increased the functional recovery involving posterior touch sensation following axotomy of PLM neuron in adulthood. Conversely, the loss of *let-7* could bypass the regeneration block due to the loss of DAF-16, a transcriptional target of DAF-2. Similarly, the loss of *daf-2* could bypass the requirement of LIN-41, a transcriptional co-factor of the *let-7* pathway. Our analysis revealed that these two pathways synergistically control targeting of the regenerating axon to the ventral nerve cord, which leads to functional recovery. The computational analysis of the gene expression data revealed a large number of genes, their interacting modules, and hub genes under *let-7* and *IIs* pathway are exclusive in nature. Our study highlights a potential to promote neurite regeneration by harnessing the independent gene expression program involving the *let-7* and Insulin signalling pathways.

## 1. Introduction

Nervous system injury leads to a wide range of behavioural consequences such as loss of sensation or fatal paralysis. Neurite regeneration from the injured stump and subsequent growth towards the target tissues often can lead to functional recovery (He and Jin 2016; Laha et al. 2017). For example, *C. elegans* sensory and motor neurons after axotomy can reach the target tissues, leading to a recovery in sensory and motor function (El Bejjani and Hammarlund 2012; Yanik et al. 2004). Similarly, in fish, the swimming behaviour is restored as the axon regrows after spinal cord injury (Becker et al. 1997; Rasmussen and Sagasti 2017). Axon regeneration potential in an organism declines with age (Basu et al. 2017; Geoffroy et al. 2016; Verdu et al. 1995) for multiple reasons (Fawcett 2020). In adulthood, injured axons face multiple challenges for correct navigation of the regenerating growth cone through the adult microenvironment (Benowitz and Popovich 2011; Brosius Lutz and Barres 2014; Fawcett 2020; Fitch and Silver 2008). Also, the cell-intrinsic capacity of the neurons to regenerate diminishes with age (Verdu et al. 1995). Therefore, functional recovery is limited in adulthood. Manipulation of intrinsic pathways and increasing neural activity together showed some promising effects in the functional restoration of retinal ganglion cells (Bei et al. 2016; Lim et al. 2016). It seems manipulation of multiple signalling cascades and transcriptional activities is what improves axon regeneration potential in adulthood (Cheng et al. 2022; Jacobi et al. 2022). For example, co-deletion of pten and sox3 promoted axon regeneration and functional recovery after the injury of RGC (Bei et al. 2016; Sun et al. 2011). Therefore, finding out synergistic controls on axon regeneration ability in the context of functional recovery is desirable.

Experiments using model organisms have been useful in deciphering the molecular mechanisms underlying the neuronal response to injury (He and Jin 2016; Richardson and Shen 2019). The molecular players essential for the regrowth of injured axons have been identified in the past decade using worm, fly and fish as model systems (Blanquie and Bradke 2018; Fawcett and Verhaagen 2018; Mahar and Cavalli 2018; Richardson and Shen 2019). *Caenorhabditis elegans* has been established as an efficient model for studying the mechanism of axonal regeneration (He and Jin 2016). Touch receptor neurons show robust regeneration response following laser axotomy at the late larval stage (Ghosh-Roy et al. 2010; Wu et al. 2007). 50% of the injured axon can fuse to their distal counterpart utilizing the phagocytosis and fusion machinery (Ghosh-Roy et al. 2010; Neumann et al. 2011). The fusion events are highly correlated with rapid functional recovery after axotomy (Abay et al. 2017; Basu et al. 2017). In later time points following axotomy, the injured axons that are targeted to the ventral nerve cords are likely to contribute to functional recovery (Basu et al. 2021). Axon regeneration and subsequent functional recovery get reduced as the worm ages (Basu et al. 2021; Basu et al. 2017; Byrne et al. 2014). It is seen that manipulation of either *let-7* miRNA or Insulin signalling (IIS) can help overcome the regeneration barriers in adulthood (Basu et al. 2021; Basu et al. 2017; Byrne et al. 2014; Zou et al. 2013). Similarly, a physical exercise involving swimming can promote functional restoration through axon regrowth following the axotomy of PLM neuron (Kumar et al. 2021).

Both *let-7* miRNA and Insulin signalling (IIS) controls the axon regeneration process in vertebrate model systems (Wang et al. 2019; Wang et al. 2018). Also, *let-7* miRNA and Insulin signalling (IIS) control other biological processes within and outside the nervous system (Dubinsky et al. 2014). *let-7* miRNA can directly control components of insulin signalling for metabolic processes in stem cells and other cell types (Jiang 2019; Kuppusamy et al. 2015; Zhu et al. 2011). Therefore, the relationship between *let-7* and IIS in *C. elegans* axon regeneration needs to be addressed.

Here using PLM neuron as a model, we correlated the anatomical features of regrowth with functional recovery at the single neuron level. We found that the functional recovery of posterior touch sensation after the axotomy of PLM neuron is synergistically enhanced upon simultaneous downregulation of *let-7* miRNA and the gene for insulin receptor *daf-2*. We found that the targeting of the injured axon towards the ventral nerve cord is enhanced in the double mutant for *daf-2* and *let-7*. *In silico* analysis of the available gene expression data in *let-7* and *daf-2* mutants revealed that these two pathways control exclusive genes and their associated networks. However, *let-7* and Insulin signalling (IIS) regulate independent genes controlling axon growth, guidance and synaptic activities. Both our experimental and computation analysis support the idea that parallel activation of gene expression under *let-7* and Insulin signaling could be highly effective for functional rewiring of injured axon.

## 2. Results

### 2.1. Simultaneous loss of *let-7* and *daf-2* synergistically enhance functional recovery of lost posterior touch sensation due to the axotomy of PLM neuron in adulthood

Since the loss of both *let-7* and *daf-2* promotes axon regeneration in adult stages in *C elegans* (Fig.1A), (Basu et al. 2021; Basu et al. 2017; Byrne et al. 2014; Zou et al. 2013), we wanted to study the relationship between these two pathways in axon regeneration using posterior touch neuron PLM as a model. PLM neurons are two of the six touch receptor neurons responsible for the posterior gentle touch sensation in *C. elegans* (Chalfie et al. 2014; Chalfie et al. 1985). Using the gentle touch assay (Basu et al., 2017) we can measure the posterior touch response index (PTRI)(Fig.S1A). After the axotomy of the PLM neuron at one side of the worm, the posterior touch response index (PTRI) is significantly reduced (Fig. S1A) (Basu et al., 2017). In later time points following axotomy PLM neuron can regenerate and contribute to the functional recovery at the day-1 stage (Fig. S1A). However, at the day-4 stage, the capacity to regenerate and restore the lost function deteriorates significantly (Fig.S1A-B) as seen before (Basu et al. 2021; Basu et al. 2017). It has been shown that loss of *let-7* miRNA and Insulin signalling (IIs) components can overcome this age-dependent decline in axon regeneration and functional recovery (Fig. 1A-B) (Basu et al., 2017, Basu et al.,2021). We asked whether these pathways act in parallel or epistatically to control axon regeneration and functional recovery. Firstly, we found that the simultaneous downregulation of *let-7* miRNA and *daf-2,* which codes for the Insulin receptor significantly enhanced the functional recovery index at 24 hours post-axotomy as compared to the respective single mutants *let-7(0)* (Tukey’s multiple comparison p<0.01**) and *daf-2(0)* (Tukey’s multiple comparison p<0.001***) (Fig.1B). The loss of *lin-41,* which is the target for *let-7* affects axon regeneration (Fig.1C), (Basu et al. 2017; Zou et al. 2013), as the gene expression program downstream of *let-7* pathway is downregulated. We found that the loss of *daf-2* in the background of the *lin-41(0)* mutant was able to bypass the requirement of *lin-41* in functional restoration post-injury (Fig. 1C-D). The recovery index value of *lin-41(0); daf-2(0)* was significantly higher than the single mutant *lin-41(0)* (Tukey’s multiple comparison p<0.001***) (Fig.1C). Similarly, we asked whether the loss the AKT kinase which acts downstream to *daf-2* in Insulin signalling (IIs) (Fig. 1A) (Paradis and Ruvkun 1998), also can bypass the requirement for *lin-41* in functional recovery. As expected, the recovery index value in *lin-41(0); akt-1(0)* was significantly higher as compared to that in *lin-41(0)* single mutant (Fig. S1C) (Tukey’s multiple comparison p<0.001***). As the loss of DAF-16, the transcription factor in the insulin signalling pathway reduces the axon targeting and functional rewiring process (Basu et al. 2021), we asked whether an enhanced gene expression program in the *let-7* mutant could bypass the requirement of *daf-16* in axon regeneration. We observed the recovery index value in *let-7(0); daf-16(0)* double mutant was significantly higher than the *daf-16(0)* (Tukey’s multiple comparison p<0.001***) single mutant but comparable to *let-7(0)* single mutant (Tukey’s multiple comparison p>0.05 ns) both at A3 (Fig.1E) and L4 stages (Fig.1F). These results suggested that the *let-7* miRNA and insulin signalling pathways synergistically enhance functional recovery after axonal injury.

**Figure 1.**
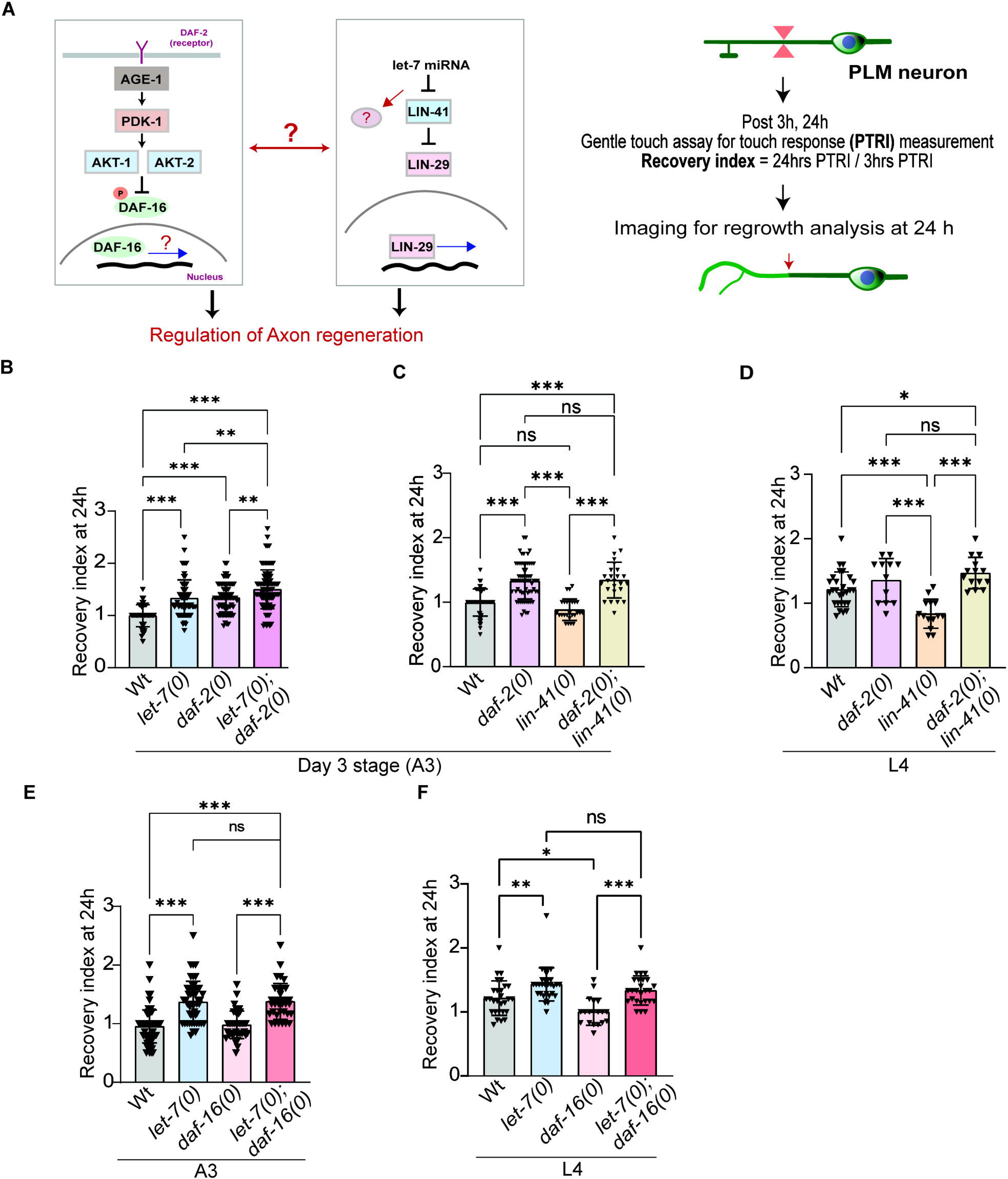
Simultaneous removal of *let-7* miRNA and insulin signalling pathways synergistically enhanced the functional recovery in day 3 adults. (A) Schematics representation of key components of let-7 miRNA signalling and Insulin signalling (left).Experimental setup for the axotomy followed by a behavioural assessment. Recovery index, the ratio between Posterior Touch Response Index (PTRI) at 24h post axotomy to 3h post axotomy, represents the functional recovery correlated to axon regeneration. Recovery index values at 24 h post axotomy are represented in the bar graph for combined genetic backgrounds (B) *let-7(0); daf-2(0)* n= number of individuals,43-95; N= Biological replicates,3-5 (C) *lin-41(0);daf-2(0)* at day 3 adulthood stage.n=25-43, N=5 and (D) *lin-41(0);daf-2(0) at* larval stage L4.n= 14-28; N= 3-5. (E) Recovery index as observed in *let-7(0); daf-16(0)* double mutant backgrounds at A3 stage.n=35-77, N=5-7 and at L4 stage (F) n= 19-31 N=4. One-way ANOVA with Tukey’s multiple comparison test was used (p<0.0001***,p<0.01**,p<0.05*).ns= non significant; Error bar represents mean ± SD.

### 2.2 The *let-7* miRNA and insulin (IIs) signalling regulates the ventral targeting events in a redundant manner

It has been shown previously that PLM neuron exhibits two broad types of regrowth patterns namely ‘fusion’ and ‘non-fusion’ events (Ghosh-Roy et al. 2010; Neumann et al. 2011) (Fig. 2A). During fusion events, the proximal and distal parts of the injured axons fuse and leads to the restoration of the lost function (Abay et al. 2017; Basu et al. 2017). When the proximal end fails to recognize its distal part and it regrows without fusion it is called a ‘non-fusion’ event. The ‘non-fusion’ class encompasses different subtypes of regrowth patterns (Fig. 2A). If the re-growing proximal part follows an anterior-posterior regrowth it is called ‘straight’. Sometimes the pattern will have multiple projections from the proximal part which is categorized as ‘multiple regrowth’ (Fig. 2A). When the re-growing end is targeted to the ventral nerve cord called ‘ventral targeting’, which also correlated with functional recovery (Basu et al. 2021). In case the regenerating axon is turned towards the dorsal side, the event is termed ‘dorsal targeting’ (Fig. 2A). We checked how these various classes of regeneration events are affected in the combination of mutants affecting *let-7* and IIs signalling. The percentage of fusion events did not increase synergistically in the *let-7(0);daf-2(0)* double muant (Fig. 2B). We observed 46% of fusion events in the *let-7(0); daf-2(0)* which was comparable to the 48% of *let-7(0)* single mutant (Chi-square test p>0.05 ns) (Fig. 2B). Addition of either *daf-2(0)* or *akt-1(0)* mutant in *lin-41(0)* background could not increase the fusion percentage significantly as compared to the *lin-41(0)* single mutant (Chi-square test p>0.05 ns) (Fig.2C, Fig.S2A). However, the % fusion events were significantly increased in the *let-7(0); daf-16(0)* background as compared to the *daf-16(0)* single mutant in both L4 and A3 stages (Fig.2D) (Chi-square test p<0.05*).This is consistent with the finding that the fusion frequency is significantly enhanced due to loss of *let-7* (Basu et al. 2017).

**Figure 2.**
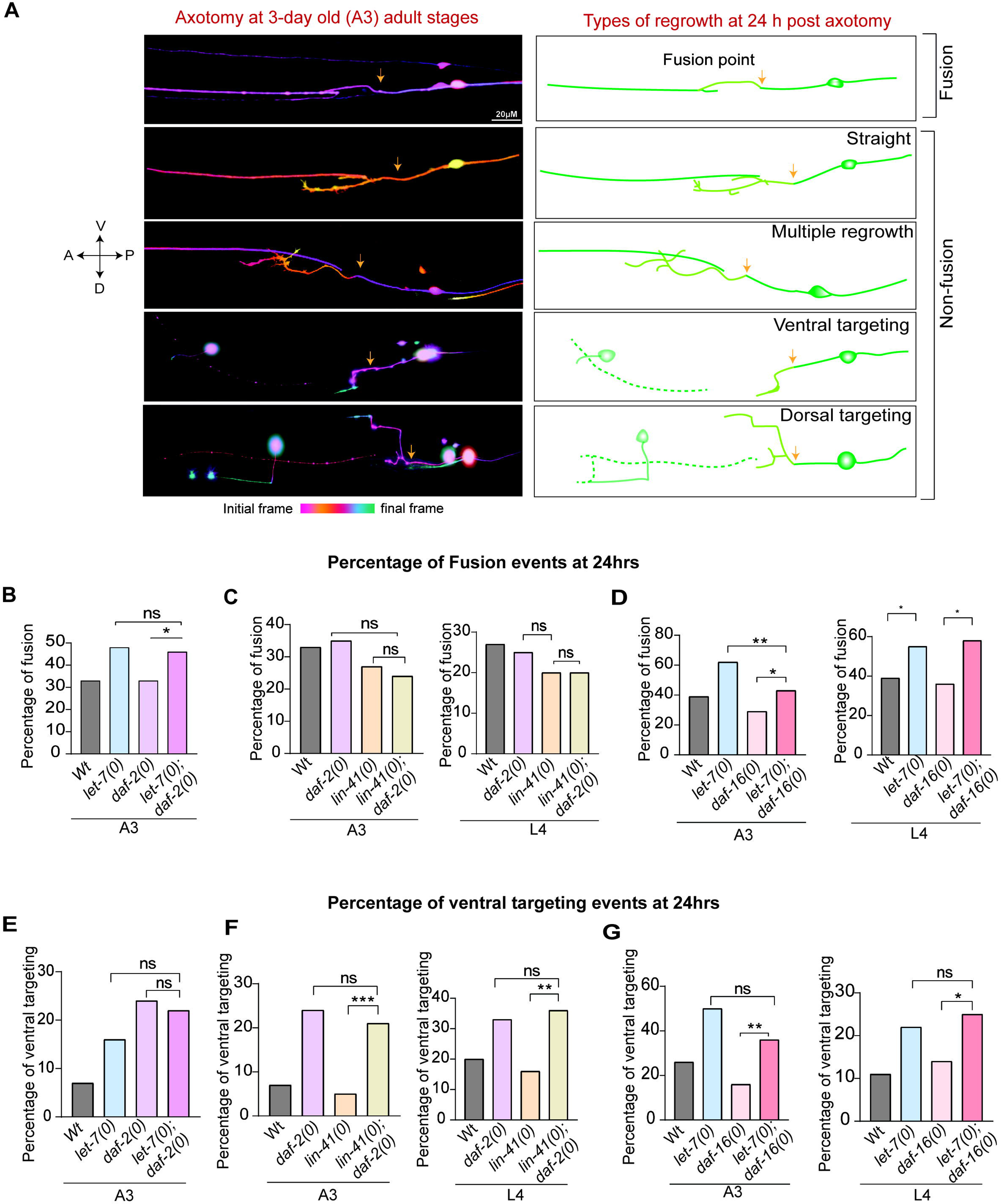
Simultaneous removal of *let-7* miRNA and insulin signalling pathways synergistically enhanced the ventral targeting events. (A) Images showing different types of regrowth observed in PLM neuron at 24 h postaxotomy. The Orange arrow represents the axotomy position. Colour coding represents the z-plane of axon regeneration. Axis represents anatomical direction (A- Anterior, P- Posterior, D-Dorsal and V-Ventral) (B) Fusion percentage at 24hrs post-axotomy in (B) *let-7(0); daf-2(0)* (C) *lin-41(0);daf-2(0)* (D) *let-7(0); daf-16(0)* at day3 adult or L4 animals as depicted.n=Number of regenerated axons; >20; N=3-5. Ventral targeting percentage at 24hrs post axotomy in combined genetic backgrounds (E) *let-7(0); daf-2(0)* (F) *lin-41(0);daf-2(0)* (G) *let-7(0); daf-16(0)* at day3 adult or L4 animals as depicted. n=Number of regenerated axons; >20;N=3-5.Chi-square test p<0.0001***,p<0.01**,p<0.05*.

Interestingly, we found that the loss of *daf-2* in the *lin-41(0)* background, enhanced the ‘ventral targeting’ events as compared to the *lin-41* mutant alone in both day3 (A3) and late larval (L4) stages (Fig. 2F). Similarly the loss of *akt-1* in *lin-41* mutant also significantly enhanced the % Ventral targeting events as compared to the *lin-41* mutant (Fig. S2B). Moreover, the loss of *let-7* in the *daf-16* background significantly elevated the percentage of ventral targeting as compared to the *daf-16* background alone in both L4 and A3 stages (Fig. 2G) (Chi-square test p<0.05* p<0.01**). On the contrary, the other classes, such as ‘straight’, ‘multi-branch’, ‘dorsal’ etc were not influenced by these two pathway in a synergistic manner (Fig. S2C-F). These observations suggested that both *let-7 miRNA* and IIS pathways might regulate the process of ventral targeting independently to enhance functional recovery.

### 2.3 Functional recovery corresponding to both the ‘fusion’ and ‘Non-Fusion’ events synergistically enhanced due to the loss of *daf-2* and *let-7*

Improved functional recovery might also be contributed partly by improved synaptic function following the regeneration events including fusion and non-fusion. To account for this possibility, we checked the functional recovery corresponding to the fusion and non-fusion events separately. Recovery index values corresponding to the fusion events showed a synergistic effect. The recovery index values of fusion events in the *let-7(0); daf-2(0)* double mutant was significantly higher than that of *daf-2(0)* single mutant (Tukey’s multiple comparison p<0.05*) (Fig.3A). In *lin-41(0);daf-2(0)* background the functional recovery in fusion events was significantly higher than the *lin-41* single mutant (Tukey’s multiple comparison p<0.05*) (Fig. 3B). Similarly, the index values of fusion events in the *let-7(0); daf-16(0)* background was significantly higher than the *daf-16(0)* mutant (Tukey’s multiple comparison p<0.001***) (Fig.3C).

**Figure 3.**
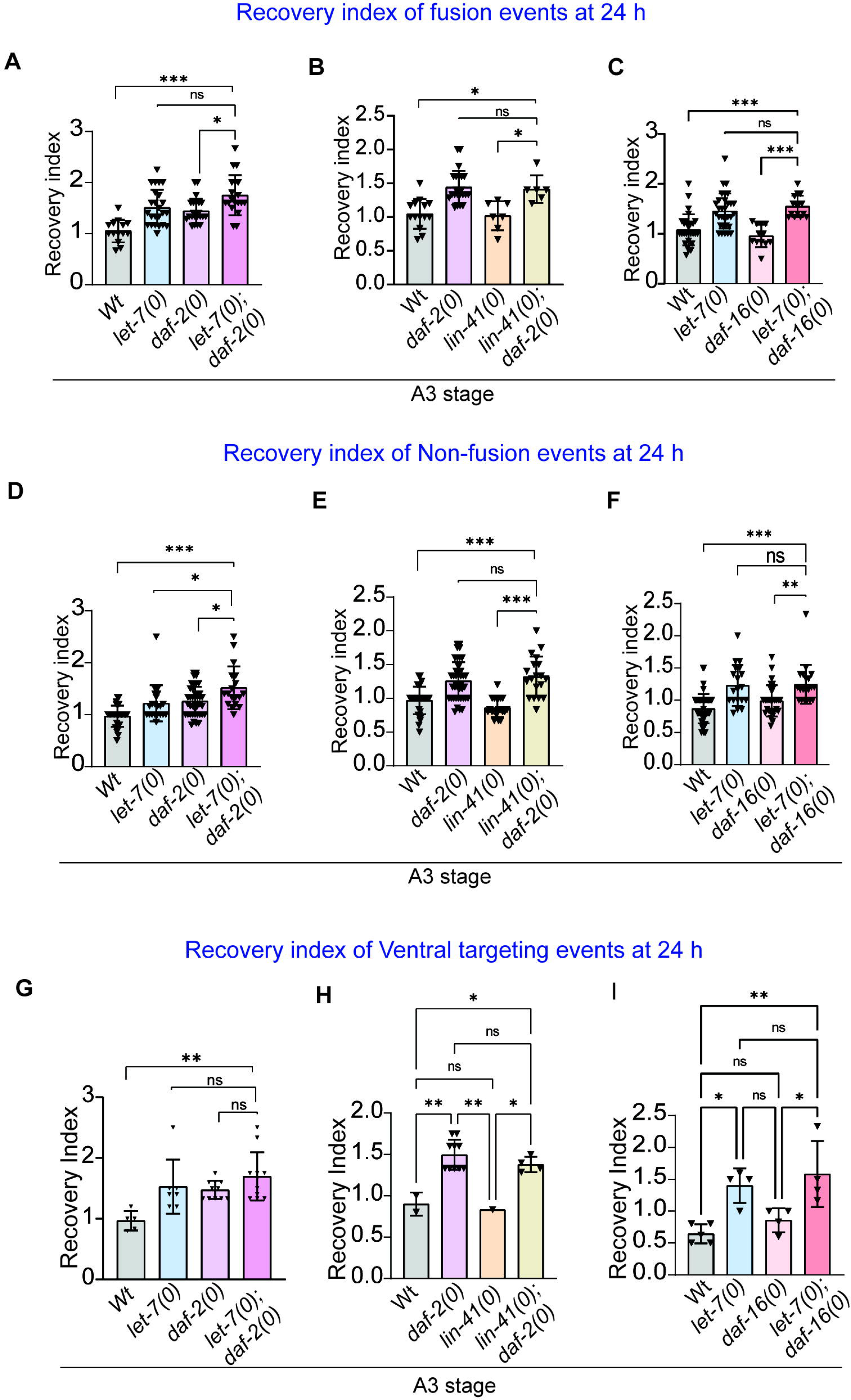
Functional recovery of non-fusion events enhanced synergistically. Recovery index values corresponding to the fusion events in combined genetic backgrounds (A) *let-7(0); daf-2(0).* n= Number of worms; 14-43.N=Biological replicates,3-4 (B) *lin-41(0);daf-2(0).* n=6-14,N=3 (C) *let-7(lf); daf-16(0).*n=; 6-9.N=12-31,N=3-4. Recovery index values corresponding to the non-fusion events in genetic combinations (D) *let-7(0); daf-2(0).*n=29-51,N=3-5 (E) *lin-41(0);daf-2(0).* n=19-42,N=3-5 (F) *let-7(0); daf-16(0).*n=19-47; N=3-4. Recovery index comparison of ventral targeting events in genetic combinations (G) *let-7(0); daf-2(0).* n=5-11,N=3-5 (H) *lin-41(0);daf-2(0).* n=1-10,N=3-5 (I) *let-7(0); daf-16(0)*.n=4-5; N=3. Axotomy experiments were done on day 3 adults. One-way ANOVA Tukey’s multiple comparison test was used (p<0.0001***,p<0.01**,p<0.05*)

The functional recovery corresponding to non-fusion events in *let-7(0); daf-2(0)* double mutant, showed a synergistic enhancement when compared to either *let-7(0)* or *daf-2(0)* single mutants (Tukey’s multiple comparison p>0.01 *) (Fig.3D). Consistent with this finding, the recovery index (RI) for non-fusion events *lin-41(0); daf-2(0)* was significantly higher than *lin-41(0)* single mutant (Tukey’s multiple comparison p>0.001***) (Fig.3E). Similarly, the recovery index (RI) corresponding to the non-fusion events in *let-7(0); daf-16(0)* mutant was significantly higher as compared to the *daf-16* single mutant (Tukey’s multiple comparison p<0.01 **) (Fig.3F).

Previous studies have shown that ventral targeting events are regulated by the DAF-16 transcription factor (Basu et al.,2021). In *daf-16(0)* mutants it has been shown that these events are non-functional. The recovery index (RI) in *daf-16(0);let-7(0)* double mutant was significantly higher as compared to the *daf-16(0)* single mutant (Fig.3I). Also the recovery index corresponding to the ventral targeting event was significantly elevated in the *lin-41(0); daf-2(0)* double mutant as compared to the *lin-41(0)* single mutant (Fig.3H). We did not observe a synergetic increase in functional recovery corresponding to other categories (multiple, straight and dorsal) in the genetic combinations (fig. S3 A-C).

These observations suggested that the functional aspect of post-regeneration events could be also synergistically regulated by the independent gene expression programs under the *let-7* and IIs pathway. This could involve genes controlling synaptic functions like synaptic maturation, and synaptic transmission.

### 2.4 IIS and *let-7* pathways control exclusive biological networks

A redundant effect of *let-7* miRNA and IIs signalling in axon regeneration could be due to the action of differential gene products regulated by these two pathways. *let-7* miRNA and Insulin signalling pathways are pivotal hubs that regulate a wide range of biological processes (Calixto et al. 2012; Chen et al. 2015; Dubinsky et al. 2014; Murphy and Hu 2013; Zhu et al. 2011). To attempt to categorize genes controlled by these two pathways in the context of axon regeneration and functional recovery, we adopted an *in silico* approach involving a computational method of network analysis (von Mering et al. 2003) of differential gene expression data. Out of the total 4697 genes significantly enriched in *let-7(0)* vs N2 microarray data (Hunter et al. 2013), 4442 genes were identified by the Wormbase and used further for analysis. Similarly, out of 9244 differently expressed genes of *daf-2(0) vs daf-2(0); daf-16(0)* RNA sequencing data (Kaletsky et al. 2016), 8589 genes are detected and used for further analysis. Between both the gene sets, only 2324 genes are common which suggests a significant non-overlapping nature of the controlled genes regulating several biological processes (Fig. 4A, Fig. S4A-D).

**Figure 4.**
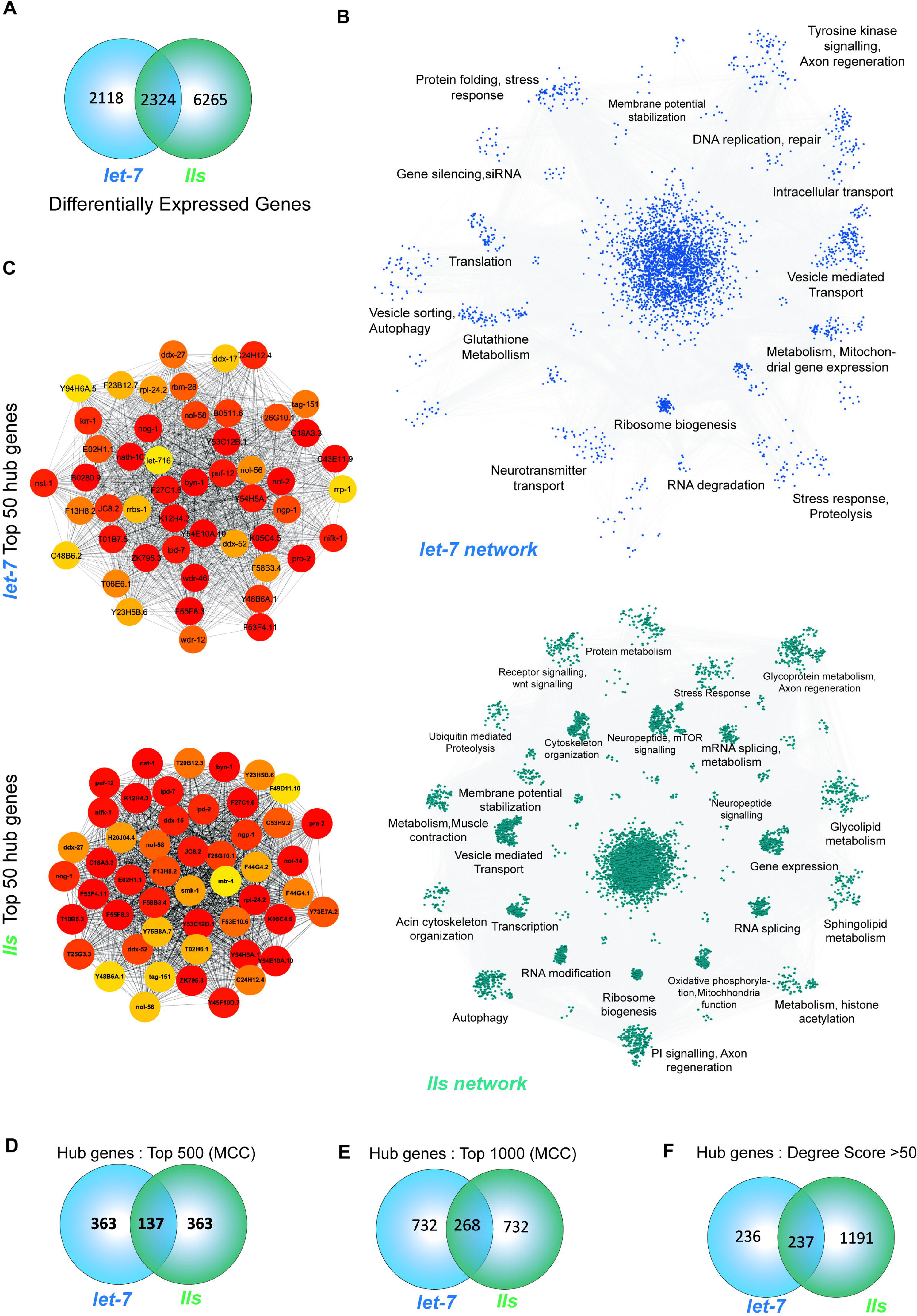
Network analysis of both *let-7* and IIs differentially expressed genes revealed the exclusivity. (A) Venn diagram showing the common and exclusivity of differentially expressed genes between *let-7* and IIS datasets. (B) Protein interaction network for *let-7 and* IIS obtained from STRING and representation of gene modules obtained by Cytoscape-MCODE analysis. (C) Top 50 hub genes predicted by Cytohubba MCC (Maximal Clique Centrality) algorithm for *let-7* and IIS dataset (D)The number of overlapping hubs among the top 500 MCC predicted hubs for *let-7* and IIS. (E) Number of Common hubs predicted as top 1000 by MCC between *let-7* and IIS data. (F) Venn diagram showing overlap status between *let-7* and IIS dataset hub genes as per the Cytohubba degree parameter (>50).

We used the STRING platform to analyse and study the interaction at the protein level (Szklarczyk et al. 2021; Szklarczyk et al. 2023). We used the STRING interface in the Cytoscape platform to obtain the protein-protein interaction network for both datasets (Shannon et al. 2003). To understand the sub-network and module in let-7 and IIS network, we further employed Molecular Complex Detection (MCODE) clustering (Bader and Hogue 2003). We found 27 and 41 modules for *let-7* and IIs-related networks respectively (Fig.4B, Table S1). To characterise the modules functionally based on their biological process and pathways, STRING-based functional enrichment was analysed in the Cytoscape. This analysis revealed that the modules under both *let-7* and IIs pathway were enriched for several important biological processes that can regulate axon regeneration and functional recovery (Fig.4B, Table S1). For example, the modules such as ‘protein translation’, ‘axon regeneration’, ‘intracellular transport’, ‘vesicular transport’ in *let-7* pathway is relevant for axon regeneration (Fig.4B). Similarly, in IIS pathway, ‘mTOR signalling’, ‘vesicle transport’, ‘cytoskeleton organization’, ‘stress response’ and many other modules are relevant for axon regeneration (Jacobi et al. 2022; Zhao et al. 2023). However, these interacting modules are categorized differentially in terms of the module functionality based on the functionally enriched nodes (protein) (Table S1). Therefore, the analysis of these module/gene network indicated that these two pathways regulate independent gene networks.

Next, we looked at the hub genes, which are highly connected in any given module/network and modulate network/module function (Barabasi and Oltvai 2004). To compare the hub genes in *let-7* and the IIS protein interaction network, we used the Cytohubba app in Cytoscape to evaluate the hub genes in 11 different algorithms. The top 50 hubs identified by the MCC (Maximal Clique Centrality) algorithm are represented in the figure shown partially overlapping (Fig.4 C, Table S2). There are only 137 and 268 common hubs in the top 500 and 1000 hubs calculated by MCC, respectively (Fig. 4 D-E, Table S2). We also screened the hub genes based on degree score which reflects the magnitude of interactions (>50) and found 473 hubs for *let-7* and 1428 hubs for the IIS network. There are only 237 hubs that are common among these two pathways (Fig.4F, Table S3).

The computational analysis revealed that the differentially expressed genes regulated by *let-7* and IIS, their interacting modules/networks (biological process), and hub genes within the interacting modules are predominantly exclusive in nature. However, these biological modules/network and hub genes could regulate growth cone initiation/elongation after axon injury (*unc-33*, *dlk-1* in IIS and *unc-33* in *let-7*), targeting (*unc-40, unc-5,vab-1,sax-3, egl-20, mom-1* in Iis and *unc-6,sax-3* in *let-7*), and synapse functionality (sad-1,*unc-104, unc-116* in Iis and *rab-3, snt-2, snb-1* in let-7), during and after neurite rewiring process.

## Discussion

The microRNA *let-7* negatively regulates heterochronic gene *lin-41* ultimately controlling larval development in *C. elegans* (Reinhart et al. 2000). This is a highly conserved microRNA and is involved in the regulation of several pathways in mammals, which also includes stem cell differentiation (Nguyen et al. 2017; Roush and Slack 2008). The previous finding shows that in the absence of *let-7* microRNA, axon regeneration is significantly enhanced in both AVM and PLM touch neurons (Basu et al. 2017; Zou et al. 2013). Enhanced axon regeneration in *let-7* promotes functional recovery as well. Both CNS and PNS neurons also show enhanced axon regeneration due to knockdown of *let-7* miRNA after optic nerve injury in mice (Wang et al. 2018). *let-7* miRNA pathway also regulates retinal regeneration (Ramachandran et al. 2010).

Similarly, insulin signalling is a key metabolic r signalling that controls several cellular processes (Haeusler et al. 2018). The absence of insulin receptor, DAF-2 correlates to improved axon regeneration as well as behavioural recovery (Basu et al. 2021; Byrne et al. 2014). In the absence of either *let-7* or *daf-2*, axon regeneration is significantly improved in aged animals(Basu et al. 2021; Basu et al. 2017). Insulin signalling plays an important role in vertebrae models of both axon and dendrite regenerations (Agostinone et al. 2018; Dupraz et al. 2013).

Our analysis highlights the combinatorial effect of these two pathways in axon regeneration and functional recovery. When the gene expression downstream of *let-7* miRNA is turned down in the *lin-41* mutant, the axon regeneration is compromised (Basu et al. 2017; Zou et al. 2013). In this study, we found that the post-injury axon regeneration and functional recovery in the *lin-41* mutant can be enhanced by boosting the gene expression involving Insulin (IIs) signalling using the *daf-2* mutant in the background of the *lin-41* mutant. Similarly, we showed that the regeneration block in the *daf-16* mutant, which compromised the gene expression of IIs signalling can be overcome by promoting the gene expression in the *let-7* cascade. More excitingly in the *daf-2;let-7* double mutant functional rewiring is synergistically enhanced. That means turning the gene expression program on in *let-7* and Insulin signalling parallelly could be a potential tool to pursue for overcoming the regeneration blocks in adulthood (Fig. 5)

**Figure 5.**
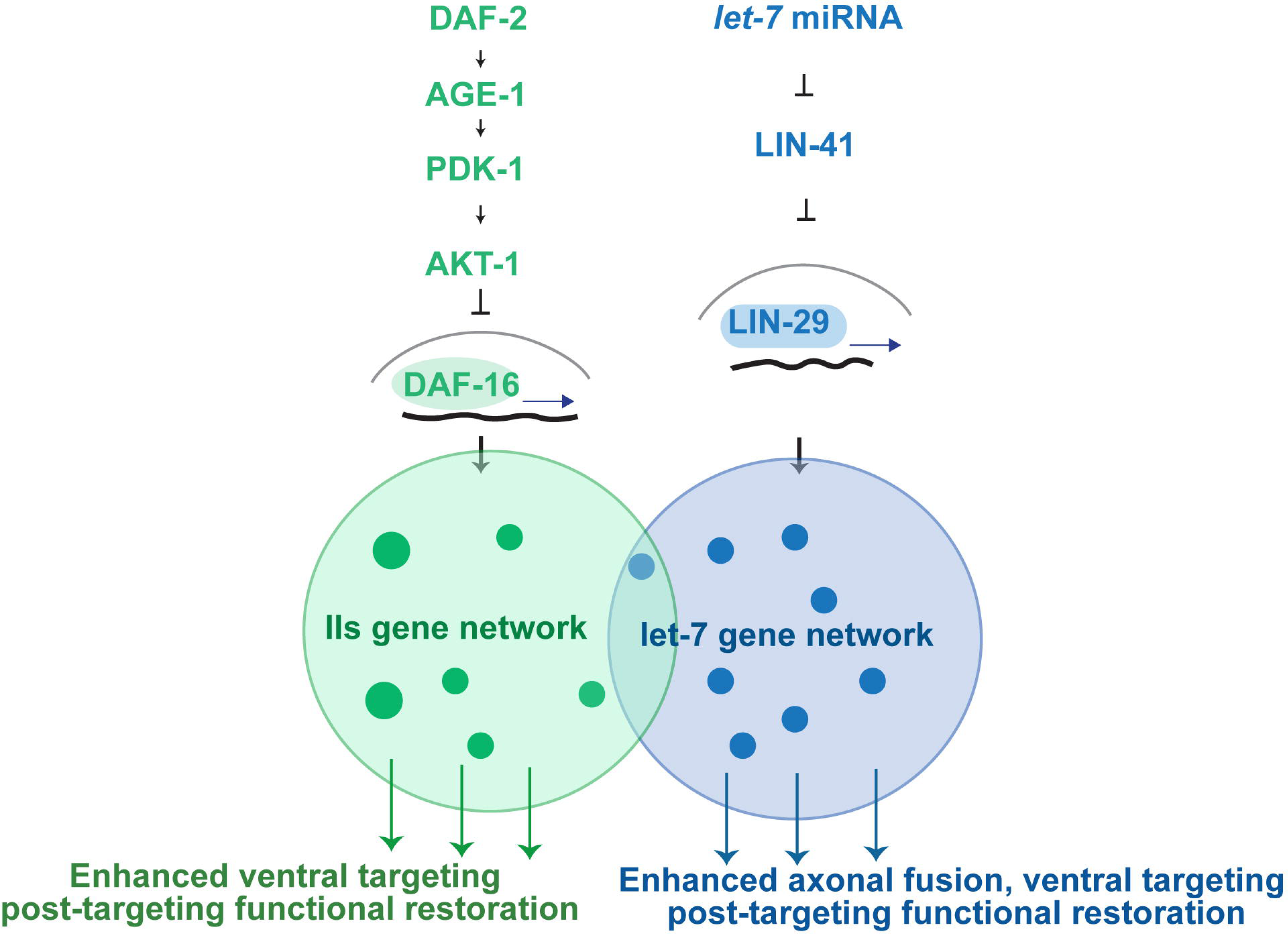
The illustration of Insulin signaling and *let-7* pathway controlling differential gene targets and their interacting modules which in turn control various aspects of axon regeneration and associated functional recovery. Filled circles within *let-7* and IIS gene network depicts the individual regulatory modules.

Successful behavioural recovery requires information transfer through circuit reestablishment. In the PLM neuron, when the regenerated axon is targeted to the ventral nerve cord region, it leads to functional recovery (Basu et al. 2021). Our analysis shows that this ventral targeting event is synergistically controlled by the *let-7* microRNA and Insulin signalling. The behavioural recovery associated to ventral targeting events also shows a synergistic effect. This suggested that the IIs and let-7 signalling controls genes related to both axon targeting as well synaptic function. Previous studies in the mammalian system report a similar role of *let-7* regulating axon guidance and synapse function (McGowan et al. 2018; Wang et al. 2019). Insulin signalling is also reported to regulate circuit establishment and synaptic function (Chiu et al. 2008; Mahoney et al. 2016; Song et al. 2003). The computational analysis shows that *let-7* and insulin signalling regulate many critical cellular processes in a non-overlapping way. The gene modules identified from the protein-protein interaction data also strongly support this. The independent regulation of several gene modules by let-7 and IIS pathway supports the finding of synergistic enhancement of functional rewiring in the genetic mutant combinations (Fig. 5). The computational analysis identified several hub genes, which can be tested further in the context of axon regeneration.

## Materials and Methods

### Genetics and *C. elegans* strains

*C. elegans* strains were grown on nematode growth medium (NGM) agar plates at 20°C following standard methods (Brenner 1974). The loss of function alleles was denoted as *’0’* throughout the manuscript except for the *daf-2* mutant. In this study, we used *daf-2 (e1368)* temperature-sensitive allele. The *daf-2(e1368)* strain was maintained at a permissive temperature of 20^0^ C. When L2 staged larvae of these mutants are transferred to the non-permissive temperature of 25°C, they produce dauer larvae (Riddle et al. 1981). This characteristic was used to make single and double mutants involving *daf-2(lf)*. For evaluating the role of DAF-2, worms were transferred to 25 °C.

All the strains used in the study are described in Table S4.

### Femtosecond lasers and axotomy

The axotomy experiments were done either at the larval stage (L4) or day 3 adult worms (A3 stage). The worms were immobilized using 0.2 μl of 0.1 μm diameter polystyrene beads (00876-15; Polysciences) in 5% agarose pads (Basu et al. 2017). Visualization of GFP labelled axon and axotomy were performed using a two-photon microscope of Bruker Corporation’s ULTIMA set-up (Basu et al. 2017). The 60X water immersion objective (NA= 1.1) was used for the imaging and axotomy of PLMs. A Laser tuned to 920 nm was used for axon visualization and a laser of 720 nm was used for severing the axon at 50 µm away from the cell body to create a gap of 7-10 μm. The cutting laser was set at the pulse width of ∼ 80 fs, irradiation pulse width of 20 ms, lateral point spread function (PSF) of ∼400 nm, and z-axis PSF of ∼1.5 μm. In all of the experiments, only one PLM was severed (either left or right). A gentle touch assay was done after a recovery duration of 3 hours and another behavioural assessment at 24 hours post-axotomy (Basu et al. 2017).

### Micropoint lasers and axotomy

Axotomy was also conducted with the use of MicroPoint lasers which is pulsed nitrogen-pumped tunable dye laser with a wavelength range of 365 nm – 656 nm. The 100x oil immersion objective (NA=1.5) was used. All the other axotomy parameters were kept the same as mentioned above.

### Gentle Touch Assay and Recovery Index

The gentle touch response assay (Basu et al. 2017; Chalfie et al. 2014; Chalfie and Sulston 1981) was performed to assess the functional correlation with axon regeneration. The posterior touch response index (PTRI) (Chalfie and Sulston 1981; Chalfie et al. 1985) was determined using the assay. According to this method, the worms were subjected to 10 alternative anterior and posterior touches with an eyelash tip and reversal of the worm was scored (Chalfie et al. 2014; Chalfie and Sulston 1981; Chalfie et al. 1985). The response was denoted as ‘1’ and no response as ‘0’ (Basu et al. 2017). Anterior touch was given to a forward-moving worm and a backward-moving worm was given a posterior touch. PTRI was assessed from both sides and later correlated to the axotomized PLM neuron side. The PTRI was calculated as the ratio of the number of responses given by individual worm per 10 stimuli at 3 hours and 24 hours post axotomy. For analyzing the extent of functional recovery we calculated the “Recovery index” which is the ratio of 24 hours PTRI and 3 hours PTRI. Therefore, a recovery index value above 1 denotes successful functional recovery.

### Anatomical axon regrowth analysis

For correlating the functional recovery with anatomical regrowth, the regenerated axon was imaged using either Nikon Ti2 or point scanning confocal microscopes after the touch assay at 24 hours. The worms were mounted on a 5% agarose pad with 10 mM Levamisole hydrochloride (Sigma L0380000) for imaging. In Nikon Ti2 the z-sections were collected using the 40x oil (NA-0.9) immersion objective and in Nikon Ti2 with the A1 Confocal system, it was done in 60x oil (NA-1.2) immersion objective. Captured z-sections were pseudo-coloured using ImageJ (Fiji®) software to represent the depth across the z-sections. Broadly the regrowth patterns were classified into ‘fusion’ and ‘non-fusion’ events (Basu et al. 2021; Basu et al. 2017; Ghosh-Roy et al. 2010). The ‘non-fusion’ events were further divided into different subtypes. The proximal neuron that regrows 35-45 μm depth towards the ventral nerve cord (VNC) was characterized as ‘ventral targeting events’ whereas the axons regrowing up to the same depth in the dorsal direction were identified as the ‘dorsal targeting’ events (Basu et al. 2021). If the proximal part regrows in the anterior-posterior direction it is called ‘straight’ and if multiple branches emanate from the proximal tip it is termed ‘multiple regrowth’.

### Quantitative analysis of axon regrowth

The optical sections corresponding to a regeneration event were z-projected together. Then the regenerated neurite length was measured using Image J (https://imagej.nih.gov/ij/download.html) (Fiji®) software and a simple neurite tracer plugin. The length of all branches from the regrowth tip was calculated for the regrowth length analysis at 24 hours.

### Experiments with the temperature-sensitive *daf-2* mutants

For the experiments using the daf-2 temperature-sensitive allele, the WT and mutant worms at the L4 stage were transferred from a permissive temperature of 20°C to a non-permissive temperature of 25°C for three days to obtain A3 stage (Day 3 old) worms for axotomy experiments. After performing axotomy, the worms were transferred back to 25°C for assessing PTRI at 3 and 24 hours.

### Analysis of differentially expressed genes

The differentially expressed genes (DEGs) for *let-7(lf)* vs N2 microarray data was obtained from (Hunter et al. 2013) and the RNA sequencing data for *daf-2(lf) vs daf-2(lf); daf-16(lf)* was used from (Kaletsky et al. 2016). Significant DEGs with p-values <0.05 were selected for further analysis. A total of 4697 genes were significantly enriched in *let-7(lf)* vs N2 background in which 4465 genes were identified by Wormbase and used for further analysis. Among these 2082 genes were upregulated and 2383 genes were downregulated. Similarly, for *daf-2(lf) vs daf-2(lf); and daf-16(lf),* 5178 genes were downregulated and 4066 genes were upregulated and further used for analysis.

### STRING database for protein-protein interaction (PPI) network

The differentially expressed genes (DEGs) were functionally characterized by the Wormcat 2.0 web tool (Holdorf et al. 2020). For protein-protein interaction network construction, we used the STRING database app in Cytoscape –v3.10.2(Shannon et al. 2003; Szklarczyk et al. 2021; Szklarczyk et al. 2023). For the prediction of interactions, we set medium confidence of 0.4 cut-off score and with 0 interactors in the full interaction mode for constructing the network. We used the MCODE tool in the Cytoscape to generate modules (Bader and Hogue 2003). The parameters for clustering were set as, Degree cutoff-2 without loops, haircut selected, Node cutoff score-0.2, K-core value 2 and maximum depth 100. The clusters having a cumulative mcode score > 4.0 were considered and the cluster was functionally annotated based on the STRING functional enrichment.

### Classification of hub genes

To identify the hub genes in the network, we used the CytoHubba tool in the Cytoscape platform which calculates the hubs in 11 different algorithms (Chin et al. 2014). Hub genes identified by MCC (Maximal Clique Centrality) and by degree score were compared. For computing feasibility, the IIS data set was trimmed with the fold change filter of ± 0.5 for the Cytohubba analysis.

### Statistics

Statistical analysis for all experiments in this article was done using Graph Pad Prism software (GraphPad Prism 9.2.1). Three or more arrays were compared using ANOVA (non-parametric) with Tukey’s multiple comparisons test. Three or more arrays were compared using ANOVA (non-parametric) with Tukey’s multiple comparisons test. Two conditions are compared with the Mann-Whitney’s t-test (non-parametric). The chi-square test was used to compare the percentage values of multiple sets in the contingency plots. For all the plots, ‘n’ (the number of samples) and ‘N’ (the number of independent replicates) are mentioned in the figure-panels and the legends, respectively. The error bars in the plots represent the standard deviation (S. D.).

## Acknowledgements

We thank NBRP, Japan and Caenorhabditis Genetics Center (CGC) for strains. CGC is supported by the NIH Office of Research Infrastructure Programs (P40 OD010440). We thank Shirshendu Dey and Swagata Dey for maintaining the 2-photon and the micro point UV laser set-up respectively. We acknowledge Atrayee Basu for sharing preliminary data and strains. This work is supported by the NBRC core fund from the Department of Biotechnology, The India Alliance DBT Wellcome Senior fellowship (Grant #IA/S/22/1/506243) to Anindya Ghosh-Roy and a grant from Science and Engineering Research Board (SERB: CRG/2019/002194) to Anindya Ghosh-Roy.

## Author Contributions

Sruthy Ravivarma, Dipanjan Roy, and Anindya Ghosh-Roy designed experiments. Sruthy Ravivarma generated mutant combination using genetic crosses, performed experiments and analyzed data. Sibaram Behera has done the in silico analysis of gene expression data. Sruthy Ravivarma, Sibaram Behera, and Anindya Ghosh-Roy wrote the manuscript.

## Abbreviations

PLM: Posterior lateral Microtubule
PTRI: Posterior Touch Response Index
RI: Recovery Index
IIS: Insulin Signaling
WT: Wild type
VNC: Ventral nerve cord

## References

Abay Z C, Wong M Y, Teoh J S, Vijayaraghavan T, Hilliard M A, and Neumann B 2017 Phosphatidylserine save-me signals drive functional recovery of severed axons in Caenorhabditis elegans. Proc Natl Acad Sci U S A 114, E10196–E10205.

Agostinone J, Alarcon-Martinez L, Gamlin C, Yu W Q, Wong R O L, and Di Polo A 2018 Insulin signalling promotes dendrite and synapse regeneration and restores circuit function after axonal injury. Brain 141, 1963–1980.

Bader G D, and Hogue C W 2003 An automated method for finding molecular complexes in large protein interaction networks. BMC Bioinformatics 4, 2.

Barabasi A L, and Oltvai Z N 2004 Network biology: understanding the cell’s functional organization. Nat Rev Genet 5, 101–113.

Basu A, Behera S, Bhardwaj S, Dey S, and Ghosh-Roy A 2021 Regulation of UNC-40/DCC and UNC-6/Netrin by DAF-16 promotes functional rewiring of the injured axon. Development 148.

Basu A, Dey S, Puri D, Das Saha N, Sabharwal V, Thyagarajan P, Srivastava P, Koushika S P, et al. 2017 let-7 miRNA controls CED-7 homotypic adhesion and EFF-1-mediated axonal self-fusion to restore touch sensation following injury. Proc Natl Acad Sci U S A 114, E10206–E10215.

Becker T, Wullimann M F, Becker C G, Bernhardt R R, and Schachner M 1997 Axonal regrowth after spinal cord transection in adult zebrafish. J Comp Neurol 377, 577–595.

Bei F, Lee H H C, Liu X, Gunner G, Jin H, Ma L, Wang C, Hou L, et al. 2016 Restoration of Visual Function by Enhancing Conduction in Regenerated Axons. Cell 164, 219–232.

Benowitz L I, and Popovich P G 2011 Inflammation and axon regeneration. Curr Opin Neurol 24, 577–583.

Blanquie O, and Bradke F 2018 Cytoskeleton dynamics in axon regeneration. Curr Opin Neurobiol 51, 60–69.

Brenner S 1974 The genetics of Caenorhabditis elegans. Genetics 77, 71–94.

Brosius Lutz A, and Barres B A 2014 Contrasting the glial response to axon injury in the central and peripheral nervous systems. Dev Cell 28, 7–17.

Byrne A B, Walradt T, Gardner K E, Hubbert A, Reinke V, and Hammarlund M 2014 Insulin/IGF1 signaling inhibits age-dependent axon regeneration. Neuron 81, 561–573.

Calixto A, Jara J S, and Court F A 2012 Diapause formation and downregulation of insulin-like signaling via DAF-16/FOXO delays axonal degeneration and neuronal loss. PLoS Genet 8, e1003141.

Chalfie M, Hart A C, Rankin C H, and Goodman M B 2014 Assaying mechanosensation. WormBook.

Chalfie M, and Sulston J 1981 Developmental genetics of the mechanosensory neurons of Caenorhabditis elegans. Dev Biol 82, 358–370.

Chalfie M, Sulston J E, White J G, Southgate E, Thomson J N, and Brenner S 1985 The neural circuit for touch sensitivity in Caenorhabditis elegans. J Neurosci 5, 956–964.

Chen A T, Guo C, Itani O A, Budaitis B G, Williams T W, Hopkins C E, McEachin R C, Pande M, et al. 2015 Longevity Genes Revealed by Integrative Analysis of Isoform-Specific daf-16/FoxO Mutants of Caenorhabditis elegans. Genetics 201, 613–629.

Cheng Y, Yin Y, Zhang A, Bernstein A M, Kawaguchi R, Gao K, Potter K, Gilbert H Y, et al. 2022 Transcription factor network analysis identifies REST/NRSF as an intrinsic regulator of CNS regeneration in mice. Nat Commun 13, 4418.

Chin C H, Chen S H, Wu H H, Ho C W, Ko M T, and Lin C Y 2014 cytoHubba: identifying hub objects and sub-networks from complex interactome. BMC Syst Biol 8 Suppl 4, S11.

Chiu S L, Chen C M, and Cline H T 2008 Insulin receptor signaling regulates synapse number, dendritic plasticity, and circuit function in vivo. Neuron 58, 708–719.

Dubinsky A N, Dastidar S G, Hsu C L, Zahra R, Djakovic S N, Duarte S, Esau C C, Spencer B, et al. 2014 Let-7 coordinately suppresses components of the amino acid sensing pathway to repress mTORC1 and induce autophagy. Cell Metab 20, 626–638.

Dupraz S, Grassi D, Karnas D, Nieto Guil A F, Hicks D, and Quiroga S 2013 The insulin-like growth factor 1 receptor is essential for axonal regeneration in adult central nervous system neurons. PLoS One 8, e54462.

El Bejjani R, and Hammarlund M 2012 Notch signaling inhibits axon regeneration. Neuron 73, 268–278.

Fawcett J W 2020 The Struggle to Make CNS Axons Regenerate: Why Has It Been so Difficult? Neurochem Res 45, 144–158.

Fawcett J W, and Verhaagen J 2018 Intrinsic Determinants of Axon Regeneration. Dev Neurobiol 78, 890–897.

Fitch M T, and Silver J 2008 CNS injury, glial scars, and inflammation: Inhibitory extracellular matrices and regeneration failure. Exp Neurol 209, 294–301.

Geoffroy C G, Hilton B J, Tetzlaff W, and Zheng B 2016 Evidence for an Age-Dependent Decline in Axon Regeneration in the Adult Mammalian Central Nervous System. Cell Rep 15, 238–246.

Ghosh-Roy A, Wu Z, Goncharov A, Jin Y, and Chisholm A D 2010 Calcium and cyclic AMP promote axonal regeneration in Caenorhabditis elegans and require DLK-1 kinase. J Neurosci 30, 3175–3183.

Haeusler R A, McGraw T E, and Accili D 2018 Biochemical and cellular properties of insulin receptor signalling. Nat Rev Mol Cell Biol 19, 31–44.

He Z, and Jin Y 2016 Intrinsic Control of Axon Regeneration. Neuron 90, 437–451.

Holdorf A D, Higgins D P, Hart A C, Boag P R, Pazour G J, Walhout A J M, and Walker A K 2020 WormCat: An Online Tool for Annotation and Visualization of Caenorhabditis elegans Genome-Scale Data. Genetics 214, 279–294.

Hunter S E, Finnegan E F, Zisoulis D G, Lovci M T, Melnik-Martinez K V, Yeo G W, and Pasquinelli A E 2013 Functional genomic analysis of the let-7 regulatory network in Caenorhabditis elegans. PLoS Genet 9, e1003353.

Jacobi A, Tran N M, Yan W, Benhar I, Tian F, Schaffer R, He Z, and Sanes J R 2022 Overlapping transcriptional programs promote survival and axonal regeneration of injured retinal ganglion cells. Neuron 110, 2625–2645 e2627.

Jiang S 2019 A Regulator of Metabolic Reprogramming: MicroRNA Let-7. Transl Oncol 12, 1005–1013.

Kaletsky R, Lakhina V, Arey R, Williams A, Landis J, Ashraf J, and Murphy C T 2016 The C. elegans adult neuronal IIS/FOXO transcriptome reveals adult phenotype regulators. Nature 529, 92–96.

Kumar S, Behera S, Basu A, Dey S, and Ghosh-Roy A 2021 Swimming Exercise Promotes Post-injury Axon Regeneration and Functional Restoration through AMPK. eNeuro 8.

Kuppusamy K T, Jones D C, Sperber H, Madan A, Fischer K A, Rodriguez M L, Pabon L, Zhu W Z, et al. 2015 Let-7 family of microRNA is required for maturation and adult-like metabolism in stem cell-derived cardiomyocytes. Proc Natl Acad Sci U S A 112, E2785–2794.

Laha B, Stafford B K, and Huberman A D 2017 Regenerating optic pathways from the eye to the brain. Science 356, 1031–1034.

Lim J H, Stafford B K, Nguyen P L, Lien B V, Wang C, Zukor K, He Z, and Huberman A D 2016 Neural activity promotes long-distance, target-specific regeneration of adult retinal axons. Nat Neurosci 19, 1073–1084.

Mahar M, and Cavalli V 2018 Intrinsic mechanisms of neuronal axon regeneration. Nat Rev Neurosci 19, 323–337.

Mahoney R E, Azpurua J, and Eaton B A 2016 Insulin signaling controls neurotransmission via the 4eBP-dependent modification of the exocytotic machinery. Elife 5.

McGowan H, Mirabella V R, Hamod A, Karakhanyan A, Mlynaryk N, Moore J C, Tischfield J A, Hart R P, et al. 2018 hsa-let-7c miRNA Regulates Synaptic and Neuronal Function in Human Neurons. Front Synaptic Neurosci 10, 19.

Murphy C T, and Hu P J 2013 Insulin/insulin-like growth factor signaling in C. elegans. WormBook, 1–43.

Neumann B, Nguyen K C, Hall D H, Ben-Yakar A, and Hilliard M A 2011 Axonal regeneration proceeds through specific axonal fusion in transected C. elegans neurons. Dev Dyn 240, 1365–1372.

Nguyen D T T, Richter D, Michel G, Mitschka S, Kolanus W, Cuevas E, and Wulczyn F G 2017 The ubiquitin ligase LIN41/TRIM71 targets p53 to antagonize cell death and differentiation pathways during stem cell differentiation. Cell Death Differ 24, 1063–1078.

Paradis S, and Ruvkun G 1998 Caenorhabditis elegans Akt/PKB transduces insulin receptor-like signals from AGE-1 PI3 kinase to the DAF-16 transcription factor. Genes Dev 12, 2488–2498.

Ramachandran R, Fausett B V, and Goldman D 2010 Ascl1a regulates Muller glia dedifferentiation and retinal regeneration through a Lin-28-dependent, let-7 microRNA signalling pathway. Nat Cell Biol 12, 1101–1107.

Rasmussen J P, and Sagasti A 2017 Learning to swim, again: Axon regeneration in fish. Exp Neurol 287, 318–330.

Reinhart B J, Slack F J, Basson M, Pasquinelli A E, Bettinger J C, Rougvie A E, Horvitz H R, and Ruvkun G 2000 The 21-nucleotide let-7 RNA regulates developmental timing in Caenorhabditis elegans. Nature 403, 901–906.

Richardson C E, and Shen K 2019 Neurite Development and Repair in Worms and Flies. Annu Rev Neurosci 42, 209–226.

Riddle D L, Swanson M M, and Albert P S 1981 Interacting genes in nematode dauer larva formation. Nature 290, 668–671.

Roush S, and Slack F J 2008 The let-7 family of microRNAs. Trends Cell Biol 18, 505–516.

Shannon P, Markiel A, Ozier O, Baliga N S, Wang J T, Ramage D, Amin N, Schwikowski B, et al. 2003 Cytoscape: a software environment for integrated models of biomolecular interaction networks. Genome Res 13, 2498–2504.

Song J, Wu L, Chen Z, Kohanski R A, and Pick L 2003 Axons guided by insulin receptor in Drosophila visual system. Science 300, 502–505.

Sun F, Park K K, Belin S, Wang D, Lu T, Chen G, Zhang K, Yeung C, et al. 2011 Sustained axon regeneration induced by co-deletion of PTEN and SOCS3. Nature 480, 372–375.

Szklarczyk D, Gable A L, Nastou K C, Lyon D, Kirsch R, Pyysalo S, Doncheva N T, Legeay M, et al. 2021 The STRING database in 2021: customizable protein-protein networks, and functional characterization of user-uploaded gene/measurement sets. Nucleic Acids Res 49, D605–D612.

Szklarczyk D, Kirsch R, Koutrouli M, Nastou K, Mehryary F, Hachilif R, Gable A L, Fang T, et al. 2023 The STRING database in 2023: protein-protein association networks and functional enrichment analyses for any sequenced genome of interest. Nucleic Acids Res 51, D638–D646.

Verdu E, Buti M, and Navarro X 1995 The effect of aging on efferent nerve fibers regeneration in mice. Brain Res 696, 76–82.

von Mering C, Huynen M, Jaeggi D, Schmidt S, Bork P, and Snel B 2003 STRING: a database of predicted functional associations between proteins. Nucleic Acids Res 31, 258–261.

Wang X, Chen Q, Yi S, Liu Q, Zhang R, Wang P, Qian T, and Li S 2019 The microRNAs let-7 and miR-9 down-regulate the axon-guidance genes Ntn1 and Dcc during peripheral nerve regeneration. J Biol Chem 294, 3489–3500.

Wang X W, Li Q, Liu C M, Hall P A, Jiang J J, Katchis C D, Kang S, Dong B C, et al. 2018 Lin28 Signaling Supports Mammalian PNS and CNS Axon Regeneration. Cell Rep 24, 2540–2552 e2546.

Wu Z, Ghosh-Roy A, Yanik M F, Zhang J Z, Jin Y, and Chisholm A D 2007 Caenorhabditis elegans neuronal regeneration is influenced by life stage, ephrin signaling, and synaptic branching. Proc Natl Acad Sci U S A 104, 15132–15137.

Yanik M F, Cinar H, Cinar H N, Chisholm A D, Jin Y, and Ben-Yakar A 2004 Neurosurgery: functional regeneration after laser axotomy. Nature 432, 822.

Zhao P, Mondal S, Martin C, DuPlissis A, Chizari S, Ma K Y, Maiya R, Messing R O, et al. 2023 Femtosecond laser microdissection for isolation of regenerating C. elegans neurons for single-cell RNA sequencing. Nat Methods 20, 590–599.

Zhu H, Shyh-Chang N, Segre A V, Shinoda G, Shah S P, Einhorn W S, Takeuchi A, Engreitz J M, et al. 2011 The Lin28/let-7 axis regulates glucose metabolism. Cell 147, 81–94.

Zou Y, Chiu H, Zinovyeva A, Ambros V, Chuang C F, and Chang C 2013 Developmental decline in neuronal regeneration by the progressive change of two intrinsic timers. Science 340, 372–376.

