## Supplementary Figures for "Synergistic control of axon regeneration and functional recovery by *let-7* miRNA and Insulin signalling (IIs) pathways"

**Figure S1**

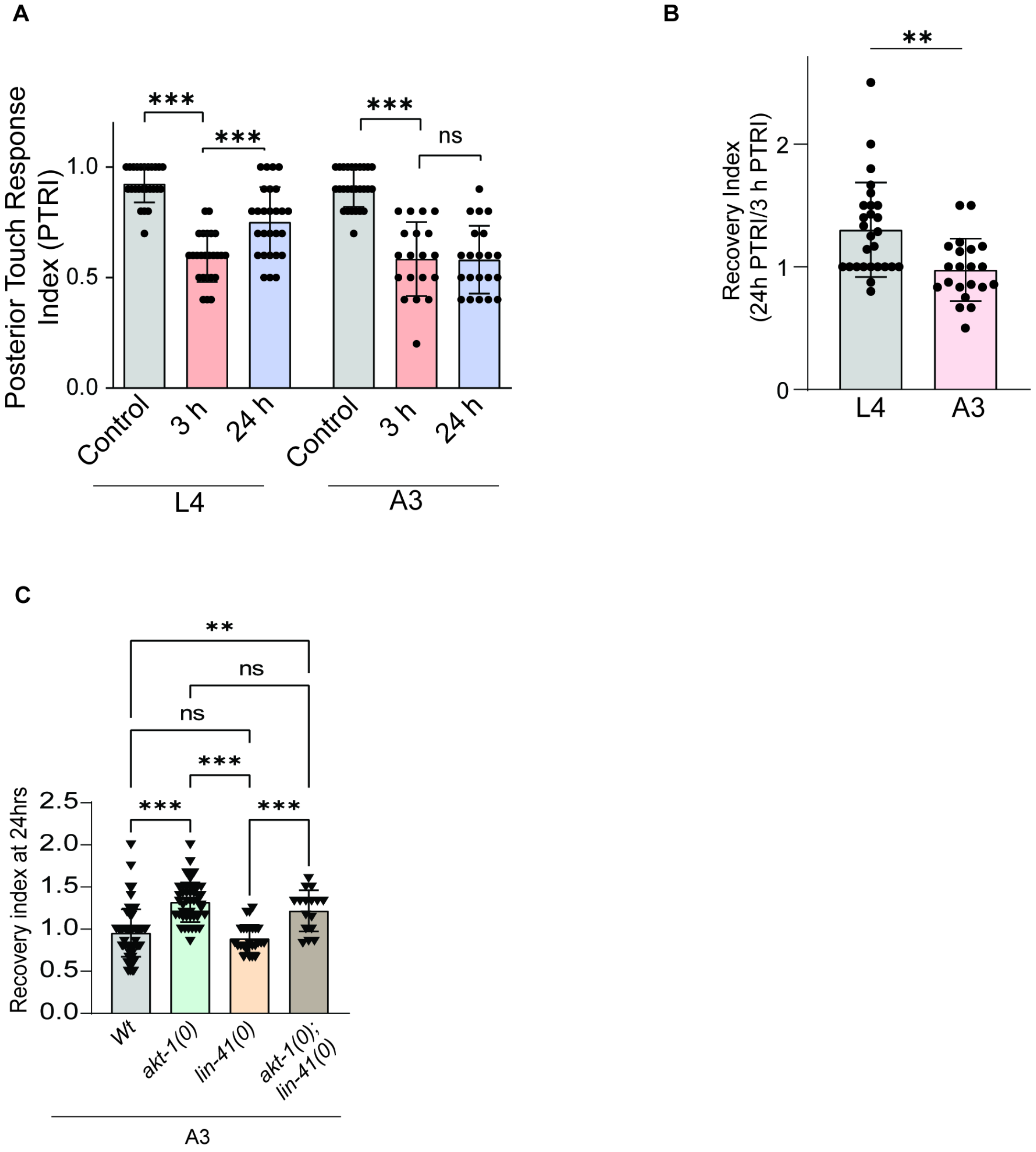


**Figure S1** (A) Posterior Touch Response Index (PTRI) values after 3h and 24 h postaxotomy in L4 stage and day 3 adult (A3) stage. n Number of worms, 22-31, N=3 (B) Recovery index is the ratio between 24h PTRI and 3h PTRI reflects reduced functional recovery at the A3 stage. n=21-27, N=3. (C) Recovery index at 24 h post axotomy in *lin-41 (0);akt-1(0)* .A3 stage axotomy. n=16-77, N=3-4. For A, B One-way ANOVA with Tukey's multiple comparison test was used (p<0.0001***,p<0.01**,p<0.05*). For B, Unpaired t-test, p<0.01**. ns= non-significant; Error bar represents mean ± SD.

**Figure S2**

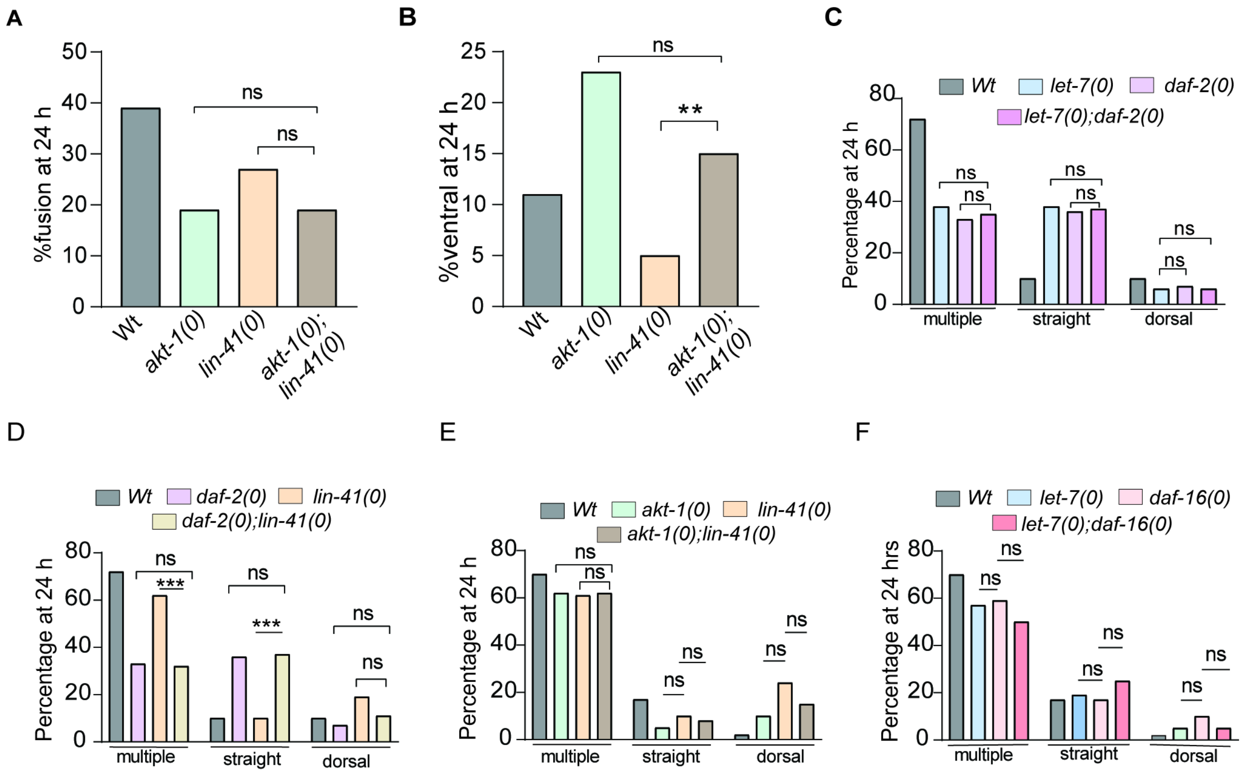


**Figure S2** (A) Percentage of fusion observed at 24h post axotomy in *lin-41(0);akt-1(0)* combination. (B) Percentage of ventral targeting observed at 24h postaxotomy in *lin-41(0);akt-1(0)* combination. Percentage of other types of non-fusion events observed in different genetic combinations (C) *let-7(0); daf-2(0)* (D) *lin-41(0);daf-2(0)* (E) *lin-41(0);akt-1(0)* (F) *let-7(0); daf-16(0)*.A3 stage axotomy and more than 20 regenerated axons evaluated. Chi-square test p<0.0001***,p<0.01**,p<0.05*. ns= non-significant

**Fig S3.**

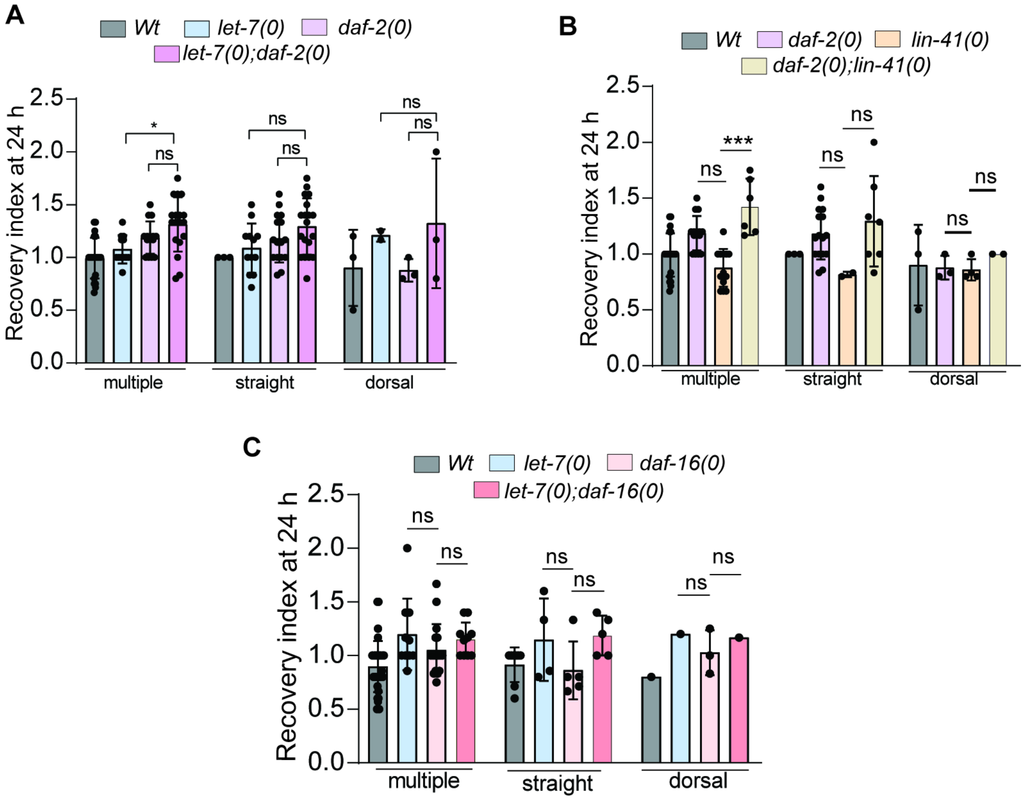


**Figure S3** Recovery index at 24 h post axotomy associated with other types of non-fusion regeneration events in different genetic combinations (A) *let-7(0); daf-2(0)* (B) *lin-41(0);daf-2(0) (C) let-7(0); daf-16(0).* A3 stage axotomy. More than 20 regenerated axons were evaluated. One-way ANOVA with Tukey’s multiple comparison test was used (p<0.0001***,p<0.01**,p<0.05*). ns= non significant; Error bar represents mean ± SD.

**Fig S4**

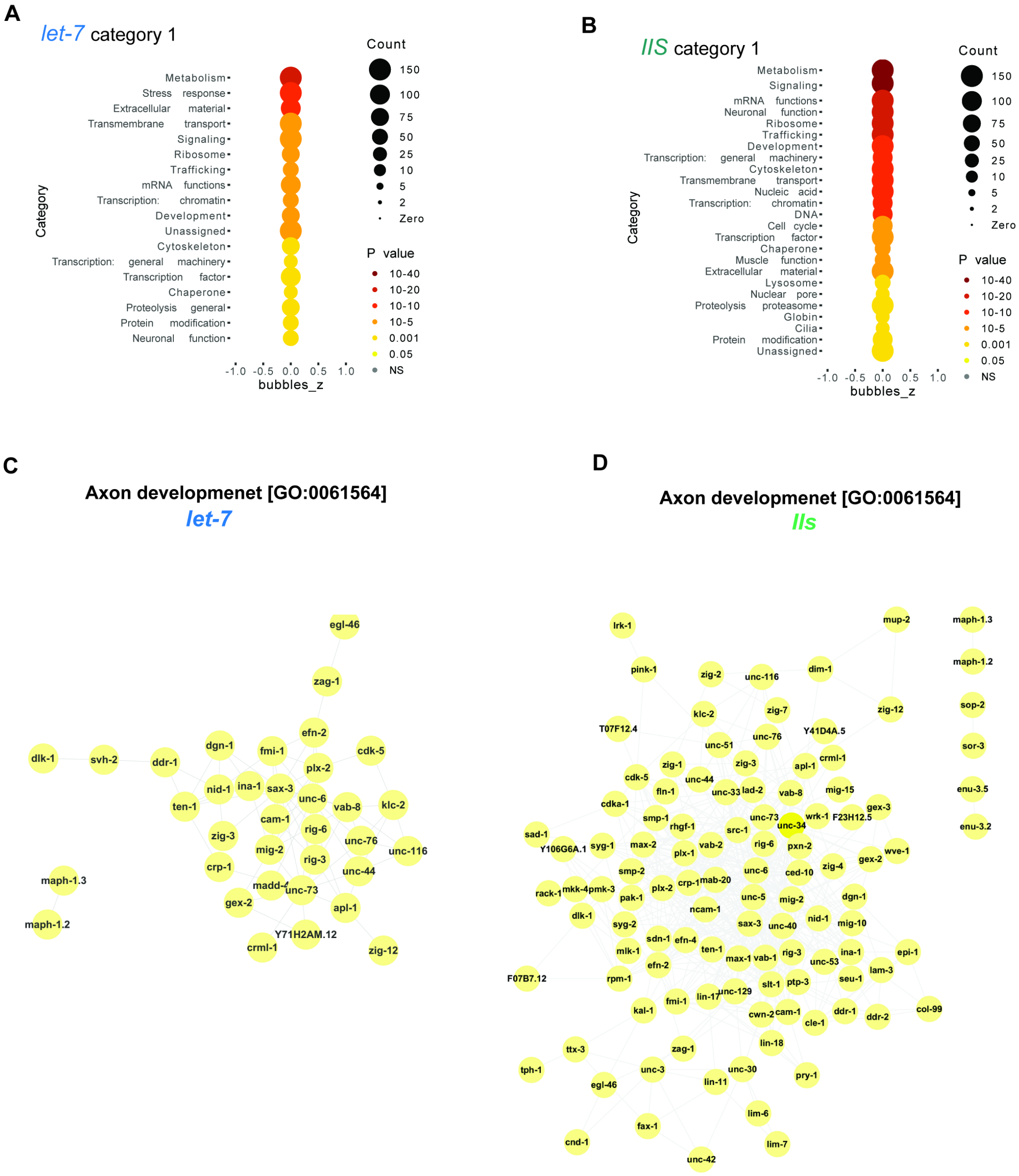


**Figure S4** Wormcat analysis results showing the broad class of (category 1) differentially expressed genes in let-7 (A) and IIS data set (B). Axon development-related genes are differentially regulated in let-7 (C) and IIS datasets (D).

**Table S1** Gene modules identified by MCODE analysis in the cytoscape platform for *let-7* dataset.

**Table S2** Top 1000 hub genes identified by CytoHubba MCC algorithm for let-7 and IIS network

**Table S3** Top hub genes identified by CytoHubba degree score (>50) for let-7 and IIS network
